# Decoding Immunomodulatory Hydrogels for Arthritis: Comparative Insights from Predictive Machine Learning and Large Language Models

**DOI:** 10.64898/2026.03.23.713755

**Authors:** Zhengkun Chen, Jie Hao, Jasmine Sarah Pye, Chenxi Zhao, Xinluan Wang, Cheng Dong, Man Ting Au, Chunyi Wen

**Author notes:** Equal contribution (Zhengkun Chen, Jie Hao).

## Abstract

Hydrogels are increasingly recognized as promising therapeutics for arthritic joints, extending their traditional role as mechanical lubricants to modulators of joint immunity. However, the rational design of these materials remains challenging, with progress largely driven by empirical experimentation. To address this, we curated a comprehensive database of 220 hydrogel formulations from 317 published studies and applied an interpretable machine learning (ML) framework to uncover the relationships between hydrogel design parameters and the arthritis severity score. Using a Random Forest algorithm, our model achieved an external validation accuracy of 0.67 in predicting effective hydrogel therapies for arthritis. Analysis revealed a clear hierarchy of design principles: the choice of functional agent, base polymer, and elastic modulus were the most influential predictors of therapeutic efficacy, with composite agents, protein-based polymers, and softer hydrogels most strongly associated with positive therapeutic outcomes. Mechanistic investigations further demonstrated that successful hydrogels promote an anti-inflammatory M2 macrophage phenotype. Benchmarking against classical statistical methods and a large language model framework showed that our ML approach provided more robust, nuanced insights into complex feature interactions. This data-driven framework offers a generalizable blueprint for the rational design of next-generation immunomodulatory hydrogels, paving the way for more effective arthritis therapies.

## Introduction

Arthritis is a prevalent and debilitating condition, characterised by chronic pain and inflammation in the joints^1^. The most common forms, osteoarthritis (OA) and rheumatoid arthritis (RA)^2^, are leading causes of disability worldwide, affecting over 8% of the global population^3,4^. The dysregulated immune microenvironment within the joint drives persistent inflammation and tissue damage^5^.

Hydrogels have emerged as a promising therapeutic platform for arthritis, as they can mimic the structural and physical properties of native articular tissue and enable sustained, site-specific treatment ^6,7^. Traditionally, hydrogel has been deployed in passive roles, such as joint lubricants^14^, drugs and cells delivery vehicles^15^, or scaffolds for tissue repair^8-10^. However, recent advances have led to the development of immunomodulatory hydrogels that actively engage the host immune response to resolve inflammation and restore joint homeostasis^11-13^. Immunomodulatory hydrogels can modulate the immune microenvironment, offering the potential for more durable and effective therapies ^16-18^.

These hydrogels may take their effect through two principal mechanisms:^19-21^: (i) serving as bioactive scaffolds that directly influence immune cell behaviour or (ii) acting as carriers for the sustained release of therapeutic agents that modulate inflammatory cascades. For example, hydrogels incorporating intrinsically bioactive materials, such as bioactive glass, can promote macrophage polarization toward the anti-inflammatory M2 phenotype^22^, while delivery-based systems can provide targeted and sustained release, such as dexamethasone to ameliorate joint inflammation and pain^23^.

Despite these advances, the rational design of immunomodulatory hydrogels remains a significant challenge. The therapeutic efficacy of these materials is governed by a complex interplay of biochemical and physicochemical properties, including composition, functional agents, and mechanical characteristics such as stiffness^24,25^. While successful applications in other fields highlight the importance of these parameters, their specific roles in arthritis therapy are not yet fully understood. The rapidly expanding body of literature, featuring diverse combinations of materials and design strategies, has rendered traditional qualitative reviews insufficient for uncovering critical design–property–outcome relationships.

To address this challenge, systematic and quantitative analysis of the literature is essential. Machine learning (ML) offers a powerful approach to identify hidden patterns and predictive relationships within large, structured datasets^26,27^. While recent advances in Large Language Models (LLMs) have enabled knowledge synthesis from unstructured text^28,29^. However, the relative strengths and limitations of these approaches for guiding biomaterial design remain unclear.

Here, we present a data-driven framework that leverages ML to systematically analyse the biomaterials literature and reveal quantitative design principles for immunomodulatory hydrogels in arthritis therapy. We curated a comprehensive dataset of 220 hydrogel formulations and their therapeutic outcome from 317 published studies. Using this resource, we trained and optimized predictive ML models to elucidate the relationship between hydrogel design features and therapeutic performance. We further benchmarked our approach against conventional statistical meta-analysis and LLM-based reasoning, providing a direct comparison of these methodologies for scientific knowledge discovery.

Collectively, our study establishes a foundational resource for data-driven arthritis biomaterials research and introduces a generalizable paradigm for AI-guided biomaterial design. By uncovering actionable design principles, our framework paves the way for the rational development of next-generation immunomodulatory hydrogels and more effective therapies for arthritis.

## Results

### Data curation and ML-based analysis workflow

A major challenge in the data-driven design of biomaterials for arthritis is the lack of comprehensive, standardized datasets^30^. Our first objective, therefore, was to construct a structured dataset connecting hydrogel design parameters to therapeutic outcomes that can be understood by the ML algorithms. For each entry, we systematically collected data across three categories: 1) hydrogel formulation and the resulting material properties, including polymer composition and encapsulated functional agents; 2) immunomodulatory effects of the hydrogels, such as cytokine secretion and cell polarization; and 3) therapeutic score, quantified by in vivo outcomes from animal models (**Fig. 1a**).

**Fig. 1.**
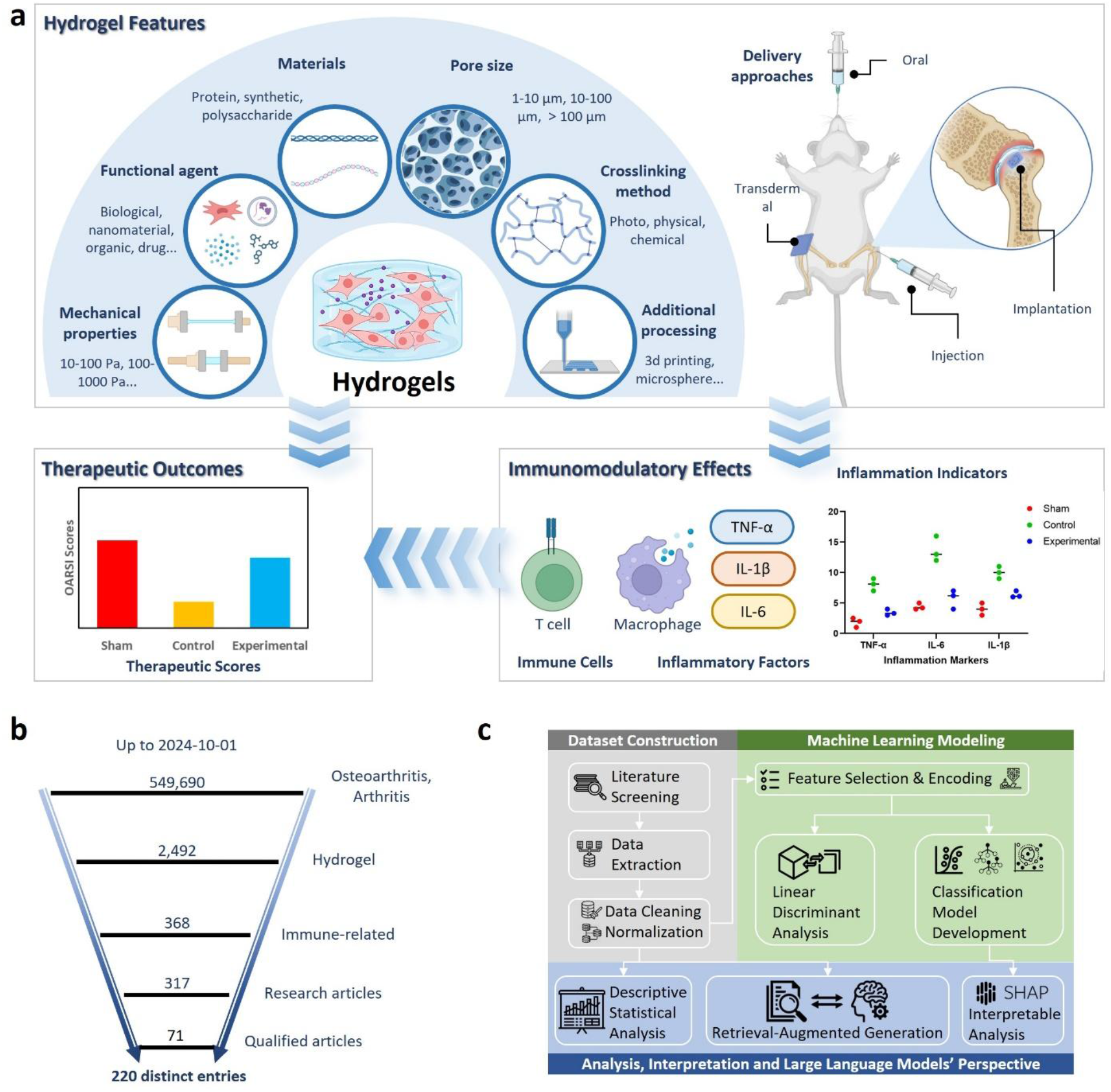
Study design and analytic framework for evaluating immunomodulatory hydrogels in arthritis. **a**, Conceptual map of the hydrogel design space analyzed in this work, including materials class (protein, synthetic and polysaccharide), pore size, crosslinking method (photo/physical/chemical), additional processing (e.g., 3D printing and microsphere formulation), mechanical properties and functional agents, alongside delivery approaches (oral, transdermal, intra-articular injection and implantation). Representative readouts include therapeutic outcomes (e.g., OARSI arthritis scores), immune-cell responses (T cells and macrophages) and inflammatory cytokines (TNF-α, IL-1β and IL-6). **b**, Procedure used for database curation using several search terms for published research articles up to 1 Oct 2024. After excluding irrelevant studies, papers without sufficient data, 71 eligible studies comprising 220 distinct entries were collected. **c**, Workflow of data construction and analysis, including literature screening, data extraction and cleaning/normalization; three distinguished approaches were used, including predictive machine learning modelling, descriptive statistical analysis, and LLM reasoning.

To build this dataset, we screened 317 research articles on using hydrogel to treat arthritis (osteoarthritis and rheumatoid arthritis) from the Web of Science. A study was included in our final dataset only if it contained complete information across all three categories, which resulted in a curated set of 220 distinct entries from 71 qualified articles (**Fig. 1b**). The curated dataset was further cleaned manually and analyzed through predictive ML algorithms, statistical analysis or LLMs (**Fig. 1c**). First, the curated data points were cleaned, normalized, and numerically encoded to create a feature matrix suitable for ML analysis. The dataset initially underwent feature-based analysis to describe overarching trends in the field’s development. Following this, we used the dataset to train a predictive model based on statistical ML algorithms to determine whether a given hydrogel formulation would be therapeutically effective. This predictive model was then interpreted using SHapley Additive exPlanations (SHAP) to identify the key design features driving its predictions. In a parallel analytical stream, we compared the insights in designing immunomodulatory hydrogels for arthritis revealed by classical statistical analysis or LLM-based reasoning against those derived from our SHAP analysis based on predictive ML.

### Polysaccharides as emerging materials for arthritis treatment

Three major material categories prevailed in the current landscape: polysaccharide-based polymers (32%), protein-based polymers (31%), and synthetic polymers (29%) (**Fig. 2a**). Among polysaccharides, three were most frequently used: hyaluronic acid (HA), a glycosaminoglycan known for its high viscosity; chitosan, a positively charged deacetylated chitin derivative; and alginate, a seaweed-derived polymer capable of rapid ionic cross-linking^31^. Protein-based materials, such as gelatin (denatured collagen), collagen (key component of the ECM), and peptides (specific cell-interactive sequences), supplied motifs crucial for tissue integration^32^. Leading synthetic polymers included polyethylene glycol (PEG), valued for its hydrophilicity and protein resistance^33^; poly (acrylic acid) (PAA), notable for its inherent stiffness^34^; and Pluronic F127, which functions as a thermoresponsive block copolymer^35^. Ultimately, material selection is driven by functional requirements^7^. Polysaccharides are prioritized for their intrinsic tissue-mimetic properties, offering excellent biocompatibility^31,36^. Proteins are highly valued for supplying vital cellular interaction motifs and bioactivity that actively guide cell response and promote successful integration^32,37^. Synthetic polymers are indispensable for achieving precise, tunable mechanical reinforcement and serving as robust cross-linking agents^33-35,38^.

**Fig. 2.**
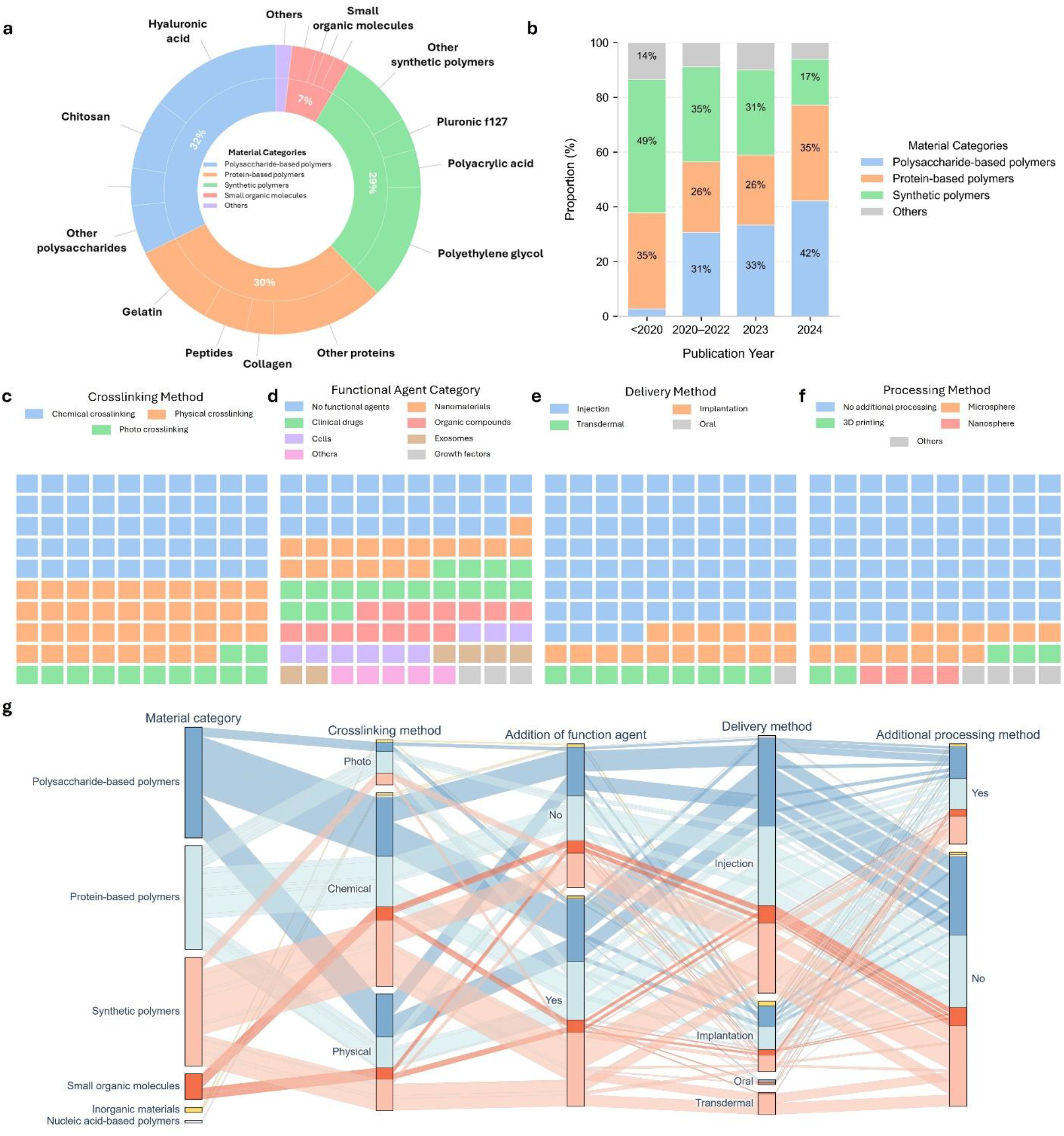
Feature-based analysis on designing immunomodulatory hydrogels for arthritis. **a**, Percentage of materials reported for hydrogel fabrication. **b**, Year-by-year shares of material categories used to fabricate hydrogels for arthritis. **c–f**, Waffle plots summarizing category proportions for crosslinking method (c), functional-agent class (d), delivery route (e), and additional processing (f). **g**, Alluvial diagram linking material category, crosslinking method, functional-agent usage, delivery route and additional processing to visualize co-occurrence patterns (colours are consistent across panels).

Notably, our analysis reveals a change in material preference over the study period from before 2020 to 2024: the prevalence of polysaccharide-based polymers increased markedly (from 3% to 43%), while the use of synthetic systems declined (from 42% to 18%) (**Fig. 2b**). This temporal shift is rooted in the deepened understanding of OA/RA pathophysiology and the maturation of chemical modification techniques ^39,40^. The marked increase in polysaccharide usage is driven by their established role in viscosupplementation (like HA) and their inherent ability to support chondrocyte viability and anti-inflammatory signaling within the joint space^41,42^. Crucially, advancements in cross-linking technologies now allow precise tuning of mechanical stability and degradation rates of these natural polymers, effectively overcoming their historical mechanical limitations^6^. Conversely, the decline in synthetic systems, despite their mechanical tunability, reflects the necessity to minimize immunological irritation and prioritize materials with intrinsic bio-recognition for successful long-term intra-articular application^43^.

Regarding crosslinking methods, chemical (49%) and physical (40%) approaches remained the most common (**Fig. 2c**). Photocrosslinking, although mechanistically a chemical method, was discussed separately in this analysis because its light-triggered mechanism could introduce distinct practical considerations^44,45^. Chemical crosslinking based on covalent bonds is the favored approach, possibly driven by the resulting mechanical robustness^6^. We noticed that the majority of the hydrogels were loaded with various types of functional agents to improve its therapeutic efficacy (**Fig. 2d**). Specifically, nanomaterials (e.g. inorganic nanoparticles and liposomes) and clinical drugs (e.g. methotrexate, and dexamethasone) were the most used agents, which enhance targeted delivery, prolong intra-articular retention, and can improve therapeutic outcomes within the joint microenvironment ^46^. Injectable formulations remained the principal route of administration (76%) (**Fig. 2e**), due to their minimally invasive delivery, conformal filling of irregular defects, and capacity for in situ gelation. Although most hydrogels were applied as bulk systems (75%), microsphere-based formulations (12%) emerged as promising strategies (**Fig. 2f**) because of their injectability, modularity, and tunable release kinetics for drug or cell delivery ^47^.

Correlation analysis among design factors revealed distinct material-strategy coupling patterns (**Fig. 2g**): synthetic polymers were more frequently associated with chemical crosslinking and often supplemented with functional agents to compensate for limited intrinsic bioactivity, whereas polysaccharide-based systems adopted either chemical or physical crosslinking and could perform effectively with or without additional functional agents, offering a broader and more versatile design space^7,43^. Similarly, protein-based polymers also demonstrated a balance between chemical and physical crosslinking and varied use of functional agents, but they exhibited a distinctly higher proportion in photocrosslinking, which is likely attributable to the high frequency of the commonly used hydrogel formulation, GelMA^48^.

### Key targets of the hydrogel-mediated immune response

Our analysis revealed distinct correlations between hydrogel material composition, their physical properties and their intended disease target. Hydrogel formulations targeting RA consistently exhibited significantly lower elastic modulus compared to those designed for OA. This observation was even more prominent in natural and composite materials **(Fig. 3a)**, suggesting a tailored mechanical approach to address the distinct pathophysiology of RA with softer tissue due to swelling and inflammation^49,50^ and OA with stiffer tissue from enlargement of bone spurs and osteochondral remodelling^51,52^. Furthermore, our analysis indicates that hydrogel pore size does not correlate with the type of diseases but rather with the class of material used **(Fig. 3b)**. Hydrogels derived from natural polymers (i.e., polysaccharides and proteins) tend to have smaller pores, whereas composite hydrogels blended from both natural and synthetic materials typically exhibit larger pore sizes. This finding is counterintuitive, as one might expect the more complex polymer network of a composite material to result in a denser structure with smaller pores.

**Fig. 3.**
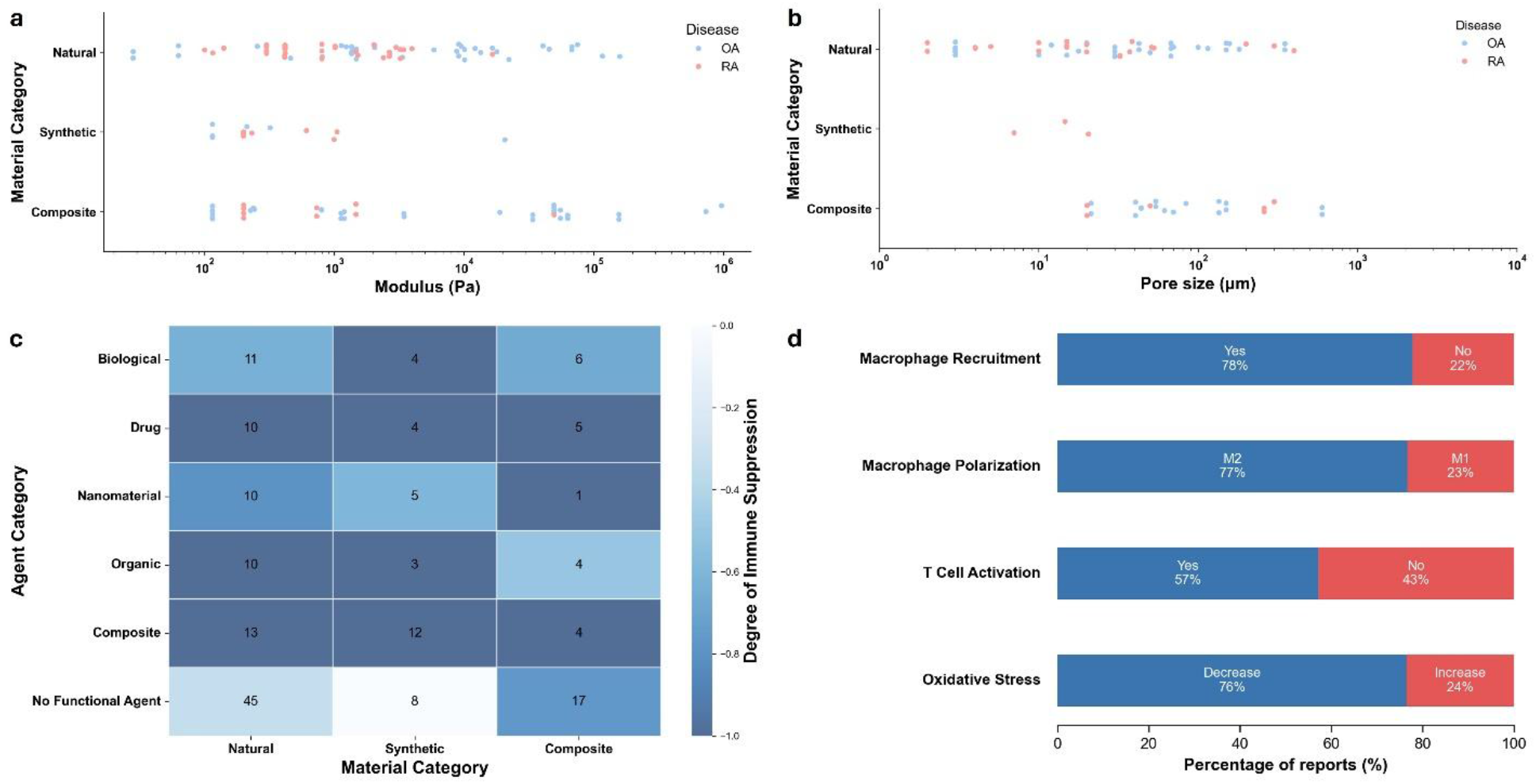
Descriptive analysis of hydrogel physical and immunological properties. **a**, Elastic modulus (Pa; log scale) by material category; each point denotes a reported hydrogel formulation (OA, blue; RA, red). **b**, Pore size (µm; log scale) by material category; each point denotes a reported formulation (OA, blue; RA, red). **c**, Heatmap of immune modulation across material categories and functional-agent classes; cells show study counts, colour indicates degree of immune suppression (lower value indicates stronger suppression; scale: -1 to 0). **d**, Proportions of studies reporting immune response after hydrogel treatment: macrophage recruitment and polarization (M2 vs M1), T-cell activation, and oxidative stress (decrease vs increase).

While all effective hydrogels aimed to suppress pathological immune responses, synthetic polymer-based hydrogels with no functional agents showed negligible immunomodulatory effects, confirming their inherent immune-inert nature. In contrast, bioactive hydrogels incorporating functional agents as the immunomodulators consistently displayed a stronger immunosuppressive effect regardless of the material used for fabrication **(Fig. 3c)**. To elucidate the immunomodulatory mechanism of the hydrogels, we analysed the reported changes in immune cells’ behavior. Successful formulations predominantly promote macrophage polarisation towards the anti-inflammatory M2 phenotype and reduce oxidative stress within the joint microenvironment. However, the impact of hydrogel design on T-cell activation remained inconclusive, with only 57% of the studies reporting a positive effect **(Fig. 3d)**. Collectively, our findings highlight that the key hydrogel material properties are more often tailored to the specific disease target. Moreover, effective immunomodulation consistently requires the incorporation of functional agents, which effectively modulate the immune response, rather than relying solely on structural properties.

### Functional agents and materials dominate the therapeutic performance

To move beyond descriptive statistics, we developed machine learning models to identify the key design parameters that govern hydrogel performance in treating arthritis. We framed this as a classification task, aiming to predict whether a given hydrogel formulation would achieve high therapeutic efficacy (defined as a therapeutic score > 0.465, **Supplementary Fig. S1**). The multicollinearity of the seven hydrogel design features was verified (**Supplementary Fig. S2**). Six commonly used classification algorithms suitable for a dataset of this size were evaluated. The Random Forest and XGBoost models demonstrated more balanced performance, with accuracy, precision, specificity, and F1-scores all exceeding 0.65 (**Fig. 4a**). These two models achieved the highest predictive power, with AUC values of 0.69 and 0.68, respectively (**Fig. 4b**).

**Fig. 4.**
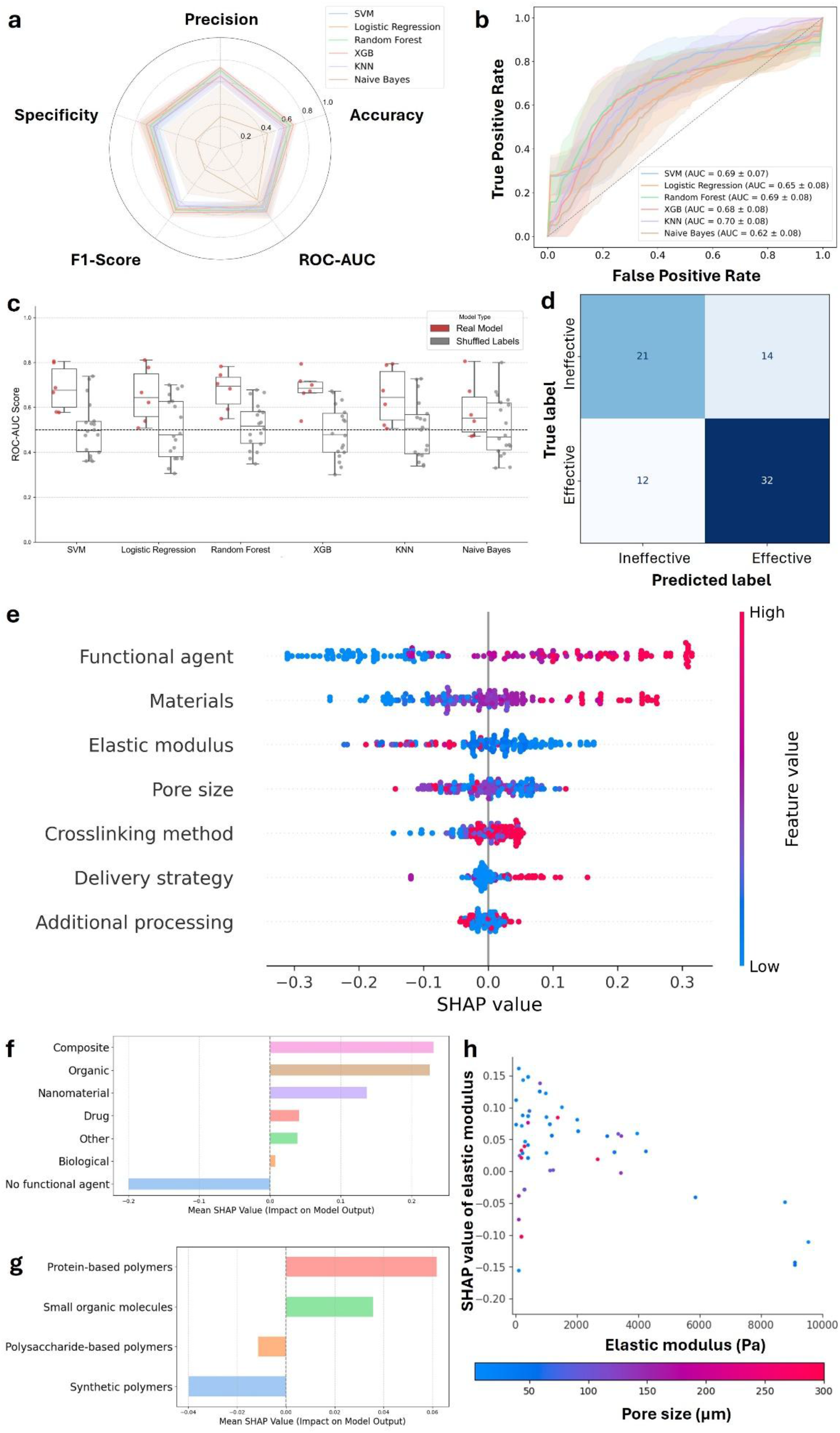
Predictive modelling and attribution of determinants of hydrogel therapeutic performance. **a**, Cross-validated performance of six classifiers for predicting whether a formulation is effective or ineffective, summarized by accuracy, precision, F1 and ROC–AUC. **b**, ROC curves (mean ± standard deviation) for each algorithm. **c**, Y-scrambling control: ROC–AUC distributions for models trained on true labels versus permuted labels (dashed line = 0.5). **d**, External validation of the Random Forest on post-October-2024 formulations. **e**, SHAP beeswarm showing global feature contributions; functional agent and material choice rank highest, followed by elastic modulus, pore size, crosslinking, delivery and additional processing. **f–g**, Category-level mean SHAP values for functional-agent classes (f) and material classes (g), indicating positive contributions from composite/agent combinations and protein-based matrices and negative contributions from no agent and synthetic polymers. **h**, SHAP dependence of elastic modulus with points coloured by pore size, showing that softer gels and smaller pores increase the predicted probability of an effective formulation.

To assess model robustness, we performed a Y-scrambling procedure across the seven algorithms. In every case, models trained on the true outcome achieved significantly higher ROC-AUC scores than those trained on permuted labels (p < 0.05), confirming that the predictive performance reflects a genuine signal rather than spurious correlation (**Fig. 4c**). In a final external validation test using unseen formulations, data from literature published after October 2024 were collected to prospectively validate the model. The Random Forest model demonstrated the best performance, achieving an overall accuracy of 0.67 (**Fig. 4d, Supplementary Fig. S3**). This indicates that the model generalizes effectively to new data and has captured the underlying design principles of successful hydrogel formulations, proving its ability to predict outcomes for novel, future hydrogel formulations for arthritis treatment.

To understand the basis for the Random Forest model’s predictions, we conducted a SHAP analysis to quantify the importance of each design feature **(Fig. 4e, Supplementary Fig. S4)**. This analysis revealed a clear hierarchy of principles that determine the effective treatment of hydrogels. The choice of functional agent was the most important predictor. The model indicated that incorporating a combination of agents (composite) was the most effective strategy for improving therapeutic efficacy **(Fig. 4f)**, with the pairing of a clinically approved drug and nanomaterials being particularly potent. We attribute this to a likely synergy between the drug’s therapeutic action and the advantages of nanomaterials as a delivery carrier. Conversely, the absence of any functional agent strongly predicted poor performance. The polymer for the hydrogel was the second most important factor. Protein-based hydrogels were strongly associated with positive outcomes **(Fig. 4g)**, while synthetic polymers were predicted to have a negative impact. Interestingly, while polysaccharides have attracted significant interest in the field, the model found them to be less beneficial, highlighting a disconnect between current research trends and the properties that yield optimal efficacy. Physical properties, elastic modulus and pore size, ranked third and fourth in importance. Dependence plots showed that hydrogel efficacy generally decreased as the elastic modulus increased, with softer gels and smaller pore sizes being most effective (**Fig. 4h**). This is likely because a soft mechanical environment facilitates cellular remodelling, while smaller pores help sustain the release of functional agents. Finally, advanced processing methods, such as 3D printing or creating injectable formulations, had a negligible impact on therapeutic outcomes in the model. This suggests their primary benefit is in improving the delivery and handling of the hydrogel rather than directly enhancing its therapeutic action.

### Suppressing inflammation benefits treatment

To identify patterns of effective hydrogel in treating arthritis through immunomodulatory and inform the design mechanism for immunomodulatory hydrogel, we conducted linear discriminant analysis to distinguish effective hydrogel formulation from non-effective ones in a 2-dimensional feature space (**Fig. 5a**). To explore the underlying mechanisms, we colour-coded the data points based on the hydrogel-induced immune response while maintaining their position in the LDA plot.

**Fig. 5.**
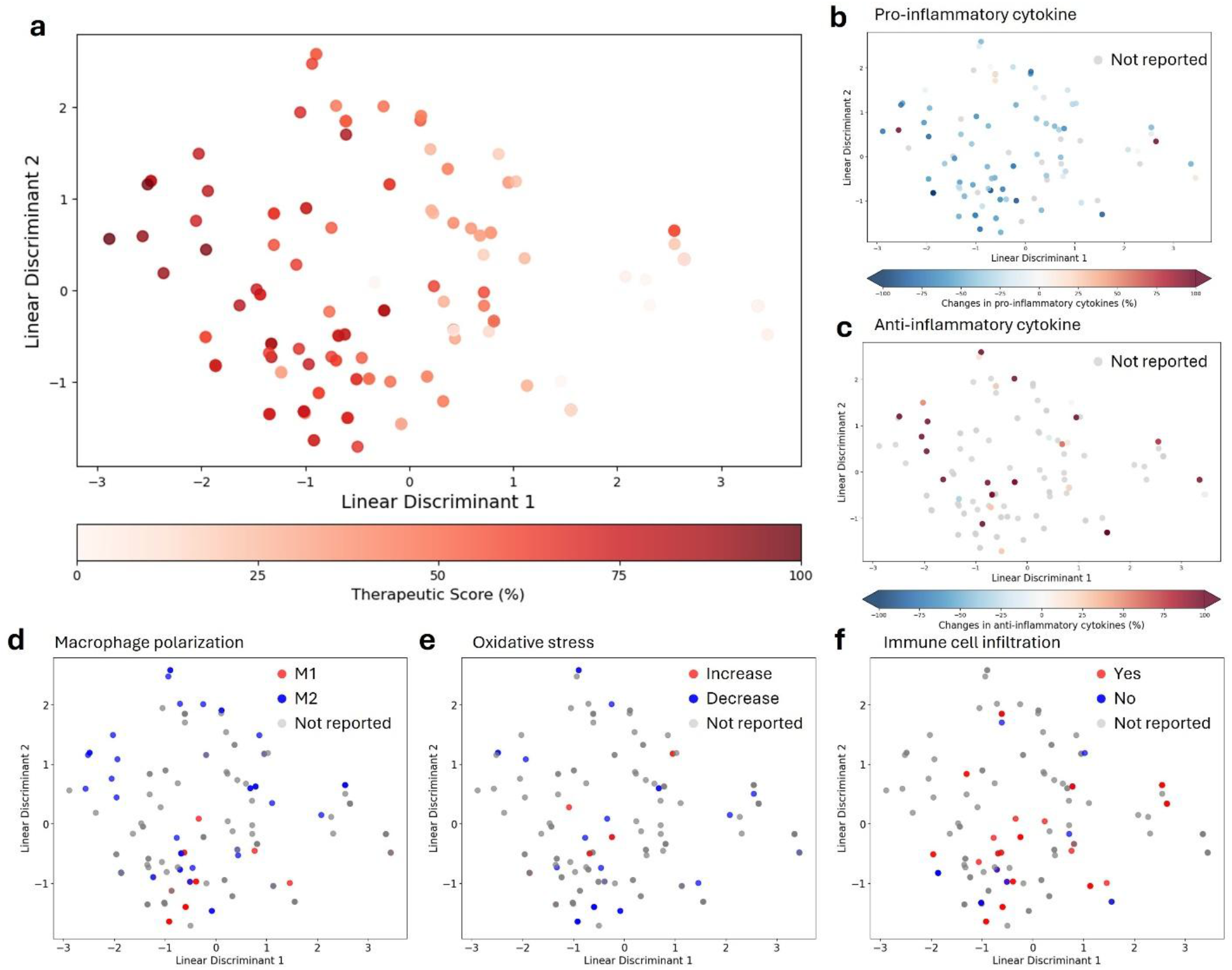
Linear discriminant analysis (LDA) of hydrogel therapeutic performance and immune response. **a**, The curated dataset projected onto two linear discriminants (LD1, LD2); colour encodes therapeutic score (%), with higher-scoring (more effective) formulations clustering on the left. **b–f**, Same LDA coordinates as in a, recoloured by immune responses to probe mechanism: changes in pro-inflammatory cytokines (b), anti-inflammatory cytokines (c), macrophage polarization (M2 vs M1) (d), oxidative stress (e), and immune-cell infiltration (f).

The analysis revealed that effective hydrogels, which cluster in areas of high therapeutic scores (darker red points in **Fig. 5a**), are strongly correlated with the suppression of inflammation. This is evident by their association with reduced pro-inflammatory cytokine release (**Fig. 5b**) and increased anti-inflammatory cytokine expression (**Fig. 5c**). This finding is mechanistically supported by the strong positive correlation between therapeutic performance and macrophage polarization (**Fig. 5d**). Specifically, hydrogel formulations that stimulated M2 polarization correlated with those with high therapeutic scores. Conversely, M1 activation was associated with poorer outcomes. The correlation with oxidative stress was ambiguous, with high therapeutic scores overlapping with both increases and decreases in oxidative stress (**Fig. 5e**), suggesting a more complex role that requires further study. Notably, while increased cell infiltration showed a moderate positive correlation with efficacy (**Fig. 5f**), its impact was less decisive than that of cytokine profiles. This is likely because cell infiltration is a necessary initial step in the immune response but is not, by itself, a guarantee of a successful therapeutic outcome.

### LLM’s perspective into the data

To benchmark the insights from our primary model, we conducted a comparative analysis of three distinct methodologies for deriving design principles from the curated dataset. Each method was evaluated on its ability to answer three fundamental questions: 1) What is the ranked importance of the main design factors? 2) What are the top three material categories? and 3) What are the top three functional agents for maximizing therapeutic efficacy? (**Fig. 6a**). Predictive ML, our primary analysis developed in previous sections, was compared with classical statistical analysis and the emerging LLM-based analysis. The classical statistical analysis employed the hypothesis-testing approach using one-way ANOVA and effect size calculations to assess the statistical significance of each design feature. The LLM-based analysis leveraged Retrieval-Augmented Generation (RAG) to synthesize insights by grounding its reasoning directly in our curated dataset as a factual knowledge base. This comparative framework allows us to assess the unique strengths and potential biases inherent in each analytical approach.

**Fig. 6.**
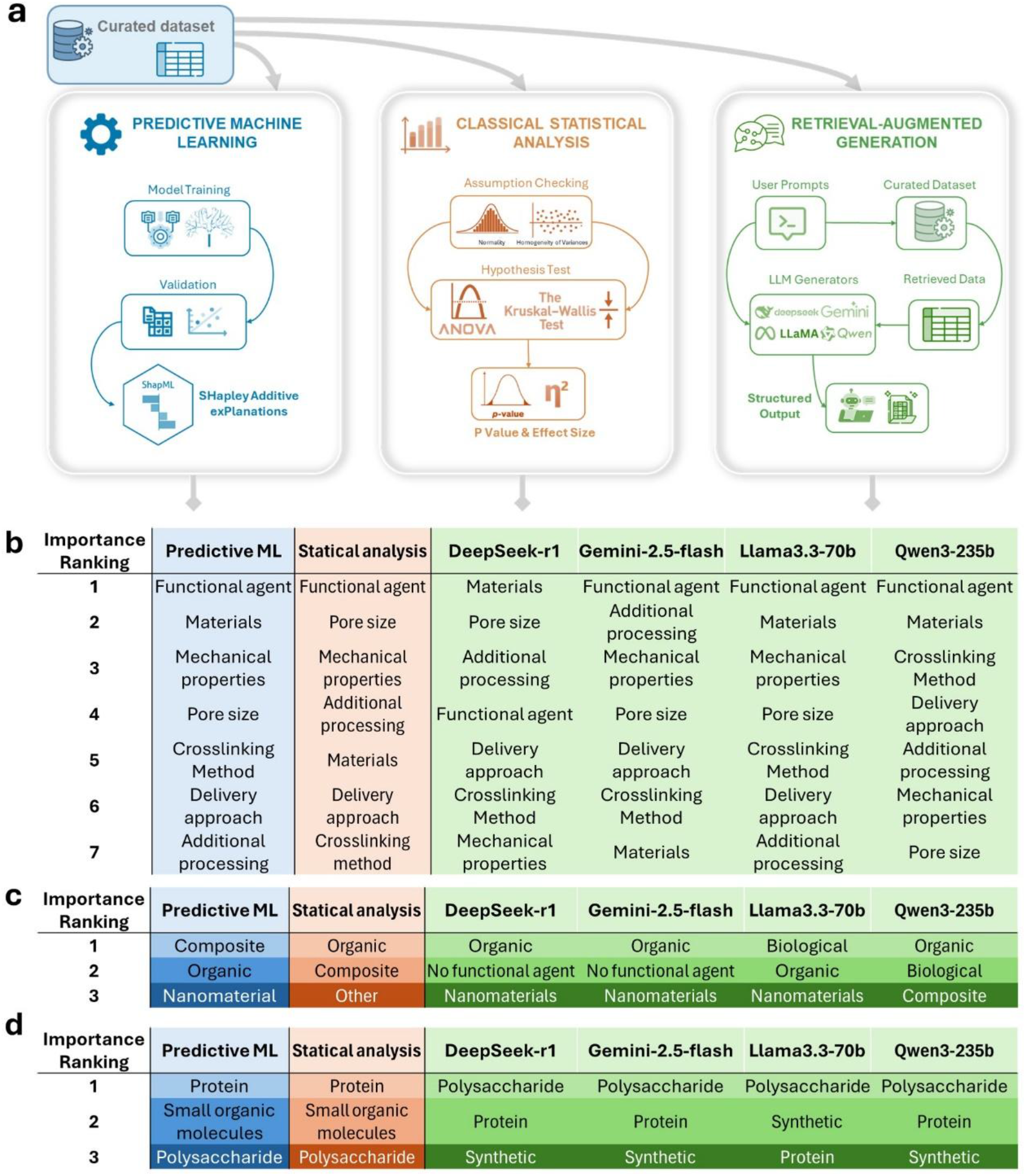
Benchmarking three approaches for deriving hydrogel design principles. **a**, Overview of the pipelines evaluated on the curated dataset: predictive machine learning (training/validation with SHAP explainability), classical statistical analysis (assumption checks, one-way ANOVA and effect sizes) and retrieval-augmented generation (RAG) with large language models grounded in the corpus. **b**, Side-by-side rankings of feature importance for therapeutic efficacy. **c**, Top three functional-agent categories by method. **d**, Top three base-material categories by method.

The three analytical methods yielded both convergent and divergent conclusions on the key principles of hydrogel design. Our predictive ML model, interpreted via SHAP analysis, identified functional agents, base materials, and elastic modulus as the top three most significant design features. In contrast, pore size, crosslinking method, and processing strategies had a negligible impact (**Fig. 6b**). The Classical Statistical Analysis agreed on the primary importance of the functional agent but ranked pore size as the second most impactful feature, with all other factors being statistically insignificant (p > 0.1). The LLM-based analyses produced more varied rankings but consistently identified the functional agent as the most critical factor (except for DeepSeek), with the Llama-3 model showing the closest overall agreement with our predictive ML model.

When examining the specific choices for functional agents, the predictive ML model identified composite, organic, and nanomaterial agents as the most effective (**Fig. 6c)**. The statistical analysis similarly favoured organic and composite agents but ranked “other” as its third choice. Strikingly, two LLMs (DeepSeek and Gemini) selected “No functional agent” as a top-three option, a counterintuitive result that likely points to a methodological bias. Regarding base materials, the predictive ML and statistical analyses shared the same ranking, favouring protein-based polymers (**Fig. 6d**). The LLMs, however, tended to rank polysaccharides as the most promising materials, followed by protein-based or synthetic materials. Overall, the results from LLMs tend to be further from the expert knowledge. Notably, the top three material and functional agent choices identified by the LLMs directly correspond to the three most frequently occurring categories in our dataset (**Supplementary Fig. S5**). This suggests that the LLM’s reasoning is susceptible to a population bias, where it tends to conflate the frequency of a category with its therapeutic efficacy.

## Discussion

Our study establishes a robust data-driven framework for the rational design of immunomodulatory hydrogels for arthritis therapy, advancing beyond the empirical approaches that have traditionally dominated the field. By systematically curating and analyzing a comprehensive dataset of hydrogel formulations, we identified quantitative design principles that underpin therapeutic efficacy in both preclinical and clinical settings.

Key findings from our analysis highlight the central importance of functional agent incorporation. Hydrogels containing composite functional agents, where multiple classes of immunomodulators act synergistically, consistently outperformed those with single agents. Material selection also emerges as a critical factor. While polysaccharide-based hydrogels are widely used due to their biocompatibility and availability^53^, our analysis indicates that protein-based hydrogels generally confer superior therapeutic benefits, likely owing to their intrinsic bioactivity and enhanced cell–matrix interactions^54^. Furthermore, mechanical and structural properties, particularly elastic modulus, were stronger predictors of efficacy than fabrication variables such as crosslinking chemistry or delivery strategy. Mechanistic insights from both the literature and our analysis converge on the induction of an anti-inflammatory, pro-reparative M2-like macrophage phenotype as a central pathway mediating the success of immunomodulatory hydrogels in arthritis^55,56^.

Despite these advances, our work also exposes critical challenges in the current biomaterials literature. A lack of standardized data reporting, especially regarding degradation kinetics, quantitative cytokine profiles, and immune cell responses, limits the reproducibility and generalizability of findings. We observed substantial heterogeneity in immune characterization, with studies often employing different cytokine panels or relying on qualitative assessments. To enable robust, cross-study comparisons and facilitate machine learning-driven insights, we advocate for the adoption of standardized immunological assays and consistent reporting of both pro- and anti-inflammatory markers (e.g., TNF-α, IL-6, CD206, IL-10). Additionally, publication bias toward positive results and subjective efficacy scoring may further distort the literature-derived data landscape.

To benchmark analytical strategies, we compared classical statistical methods, predictive ML, and LLM-based reasoning. Classical statistical analysis, relying on univariate correlations such as ANOVA, is well-suited to detecting strong direct associations but inherently incapable of capturing multivariate or conditional dependencies. For instance, the efficacy of a particular polymer may depend on its combination with a specific functional agent, a relationship invisible to univariate tests. The LLM-based reasoning approach demonstrated limitations in quantitative ranking and feature interpretation. Even when grounded with retrieval-augmented data, the LLM exhibited popularity bias^57^, over-weighting frequently mentioned terms and misinterpreting null or categorical variables, such as ranking “no functional agent” among top-performing designs. This raises concerns about the reliability of using language models to guide experimental design and materials discovery. In contrast, our predictive ML framework, particularly the tree-based ensemble models, excels at mapping nonlinear and high-dimensional feature interactions^58^, enabling a more holistic understanding of how compositional and physical parameters jointly influence therapeutic outcomes. When supported by a well-curated dataset, such models can serve as powerful tools for trend analysis, hypothesis validation, and rapid screening of candidate formulations.

In summary, our study demonstrates the potential of machine learning to extract and generalize actionable design rules from the biomaterials literature, offering a blueprint for the rational engineering of next-generation immunomodulatory hydrogels for arthritis therapy. By integrating predictive modeling and benchmarking analysis, we highlight the promise and the limitations of AI-driven biomaterial design. We envision that continued advances in data standardization, model interpretability, and integration with experimental pipelines will accelerate the transition from empirical material development to truly predictive, mechanism-informed biomaterials engineering.

## Methods

### Data curation and cleaning

The dataset was collected from articles published between January 1, 1990, and October 1, 2024, and collected by Web of Science for. The search strategy combined terms for hydrogels, specific arthritis types, and immunology: (hydrogel* [All Fields] AND osteoarthritis [All Fields] AND immun* [All Fields]) OR (hydrogel* [All Fields] AND “Rheumatoid arthritis” [All Fields] AND immun* [All Fields]). The search was limited to the document types of Article, Early Access, Proceeding Paper, or Book Chapters. The external validation dataset was compiled using the same search criteria but for papers published between October 1, 2024, and July 1, 2025. Only primary research articles that reported in vivo animal study data and explicitly discussed the immunomodulatory mechanism or resulting immune response of the hydrogel therapy were included.

For each qualifying study, relevant data were manually extracted and curated into a structured database. The collected parameters included: 1) hydrogel fabrication conditions, 2) physical properties, 3) immunomodulatory properties, 4) therapeutic performance metrics, and 5) other details such as the animal model used.

### Predictive machine learning analysis

In the complete dataset, entries without quantification on the therapeutic performance were excluded, yielding 162 qualified entries, including fabrication conditions and therapeutic performance metrics. Missing numerical features (i.e., elastic modulus and pore size) were imputed using the global mean of their respective columns. Categorical features were transformed using target encoding, with the exception of “processing usage”, which was transformed into a binary variable. The therapeutic performance, therapeutic score, was transformed into a binary variable by setting a threshold at 0.46, where any score above it is considered to be effective and below is ineffective. This preprocessing pipeline resulted in a 7-dimensional numerical description for each of the 162 hydrogel formulations. These features served as the inputs and the therapeutic score as the output for six distinct ML classification algorithms: Support Vector Machine (SVM), Logistic Regression, Random Forest, XGBoost, K-Nearest Neighbors (KNN), and Naïve Bayes. Models were evaluated using a repeated stratified five-fold cross-validation (n_splits=5, n_repeats=3, shuffle=True). Hyperparameter optimization was conducted within a nested cross-validation framework. For each training fold, an exhaustive grid search with an internal five-fold validation (cv=5) was used to identify the optimal hyperparameters, maximizing the roc_auc score. The model with the best-performing parameter settings was then evaluated on the held-out test fold. Model performance was assessed using accuracy, precision, sensitivity, F1-score, and the area under the receiver operating characteristic curve (ROC-AUC), as implemented in scikit-learn. To interpret the predictive models, SHAP (SHapley Additive exPlanations) analysis was performed. Each of the six finetuned models was trained on the complete dataset, and a model-specific explainer (e.g., TreeExplainer, LinearExplainer, KernelExplainer) was used to retrieve the SHAP values for every feature. Features were then prioritized and ranked based on their mean absolute SHAP value.

Separately, a Linear Discriminant Analysis (LDA) classifier (scikit-learn) was fitted to the complete set. The LDA was used to reduce the dimensionality of the data to a two-dimensional plane, enabling visualization and the correlation of design features with hydrogel efficacy based on the spatial positioning of the data points.

### Classical statistical analysis

To evaluate feature-level differences across three or more independent groups, we first examined statistical assumptions for each comparison. Normality of residuals was assessed using the Shapiro-Wilk test (H_0_: normally distributed residuals), supported by visual diagnostics including Q-Q plots and histograms. Homogeneity of variances was examined using Levene’s test (H_0_: equal variances across groups). Assumptions were considered satisfied when p > 0.05 for both tests. When the normality and variance homogeneity assumptions held, group differences were evaluated using One-way Analysis of Variance (ANOVA). In cases where either assumption was violated, the non-parametric Kruskal-Wallis H test was applied to compare the distributions across groups. For each statistical test, the corresponding null hypothesis, test statistic, and p-value were obtained. To estimate the magnitude of group effects, effect sizes were calculated alongside hypothesis testing. For ANOVA, Eta-squared (η^2^ = SS_effect / SS_total) was computed, while for the Kruskal-Wallis test, a ranked-data Eta-squared metric (η^2^_ranked = H / (N + 1), where N is the total sample size) was used. These effect size values were further utilized to rank the features based on their relative influence. Statistical significance was defined as p < 0.05.

### LLM-based Analysis with RAG

To assess the LLM’s capacity for scientific reasoning, we implemented a Retrieval-Augmented Generation (RAG) workflow in our curated dataset. First, a searchable knowledge base was created from our 220-entry dataset. A textual representation for each hydrogel formulation was generated by concatenating its design factors, hydrogel properties, associated immune response, and therapeutic outcome into a single string. These descriptions were then encoded into a high-dimensional vector space using Google’s text-embedding-004 model (task_type=“RETRIEVAL_DOCUMENT”), creating an embedding vector for each database entry. The analysis was executed via a query-and-retrieval workflow. A high-level analytical query, searching for the key design principles for immunomodulatory hydrogels with a high therapeutic score, was first embedded using the same text-embedding-004 model (task_type=“RETRIEVAL_QUERY”). A cosine similarity search was performed between this query vector and the entire database to retrieve the top_k (k=15) most relevant data entries. This structured data block was integrated into a comprehensive prompt that instructs the LLMs to act as a materials scientist and base their analysis exclusively on the provided data. The model was then tasked with ranking the feature importance of design factors, identifying the top materials, and identifying the top functional agents. This complete, augmented prompt was processed by four distinct models: llama-3.3-70b-instruct, deepseek-r1, and qwq-32b (via the OpenRouter API), and gemini-2.0-flash (via Google Generative AI).

## Data availability

The authors declare that all data supporting the findings of this study are available within the main text and Supplementary Information. The curated dataset was deposited in GitHub (https://github.com/wenchunyi-polyubme/ArthritisHydrogel).

## Code availability

The source code for the feature-based analysis, model training and LLM-based reasoning has been deposited in the repository ‘ArthritisHydrogel’ (https://github.com/wenchunyi-polyubme/ArthritisHydrogel).

## Acknowledgements

This work was supported by RGC GRF (15100324; 15100821); ANR/RGC JRS (A-PolyU503/24); TRS (T11-709/21-N); NFSC/RGC (N_PolyU520/20; N_PolyU542/25); ITF MHKJFS (MHP/037/23; GHP/231/22GD); RIF (R4024-23). Z.C. did not receive any of these funds.

## Author contributions

Z.C. and C.W. conceived and supervised the project. Z.C., J.H., J.S.P. and C.Z. curated and cleaned the dataset. Z.C. and J.H. performed the data analysis and data visualization. X.W., M.T.A. and C.W. contributed to data interpretation and mechanistic insight. Z.C., J.H., M.T.A., and C.W. drafted the manuscript with input from all authors. All authors discussed the results, revised the manuscript critically for important intellectual content, and approved the final version of the manuscript.

## Competing interests

The authors declare no competing interests.

## Notes

### Competing Interest Statement

The authors have declared no competing interest.

