## Supplementary Information for "Decoding Immunomodulatory Hydrogels for Arthritis: Comparative Insights from Predictive Machine Learning and Large Language Models"

* *Equal contribution*

### Data cleaning

All relevant information was manually collected from the research articles. The initial data curation included: the type of disease, material used (i.e. the category of the materials and the exact material name), material modifications, crosslinking method, functional agent (i.e. the category of the functional agent and the exact agent name), delivery method, and processing method. We also extracted physical properties, including mechanical properties (compressive modulus, tensile modulus, and storage modulus), pore size, and degradation half life. Immunological data was captured, including responded immune cells, the type of immune response experiment, changes in specific pro- and anti-inflammatory cytokines, macrophage recruitment, macrophage polarization, T cell activation, and oxidative stress. Finally, we collected data on the animal model, any surgical intervention, and all reported clinical scores (e.g., Total Mankin Score, OARSI, ICRS Score, MODS, Pineda score, Synovitis Score, RA Score).

Mechanical properties, changes in overall cytokine production, and clinical scores in this extensive database were then processed into a final set of harmonized features for analysis. To compare the clinical performance across different scoring scheme, a harmonized score, Therapeutic Score, was calculated by normalizing the experimental group's score with respect to its control group or the scoring scheme's limits. The specific normalization formulas were as follows:

- Total Mankin Score, OARSI score, synovium score, or RA score: $\textit{therapeutic score}= \frac{\textit{No treatment group-Experimental group}}{\textit{No treatment group-Sham group}}$, assuming sham group = 0 for Total Mankin Score if the score was not reported
- ICRS Score or Modified O'Driscoll histological score: $\textit{therapeutic score}= \frac{\textit{Experimental group-No treatment group}}{\textit{12- No treatment group}}$, where 12 is the maximum score in the scheme
- Pineda score: $\textit{therapeutic score}= \textit{1-} \frac{\textit{Experimental group}}{\textit{No treatment group}}$

All normalized therapeutic scores were bounded, setting any value >1 to 1 and any value < 0 to 0. To process the data for the classification models, these continuous scores were then binarized. Based on the score distribution (**Figure S1**), a threshold of 0.465 was selected. Any hydrogel formulation with a score ≥ 0.465 was classified as "effective" (n=87), while any hydrogel formulation with score < 0.465 was classified as "non-effective" (n=75).

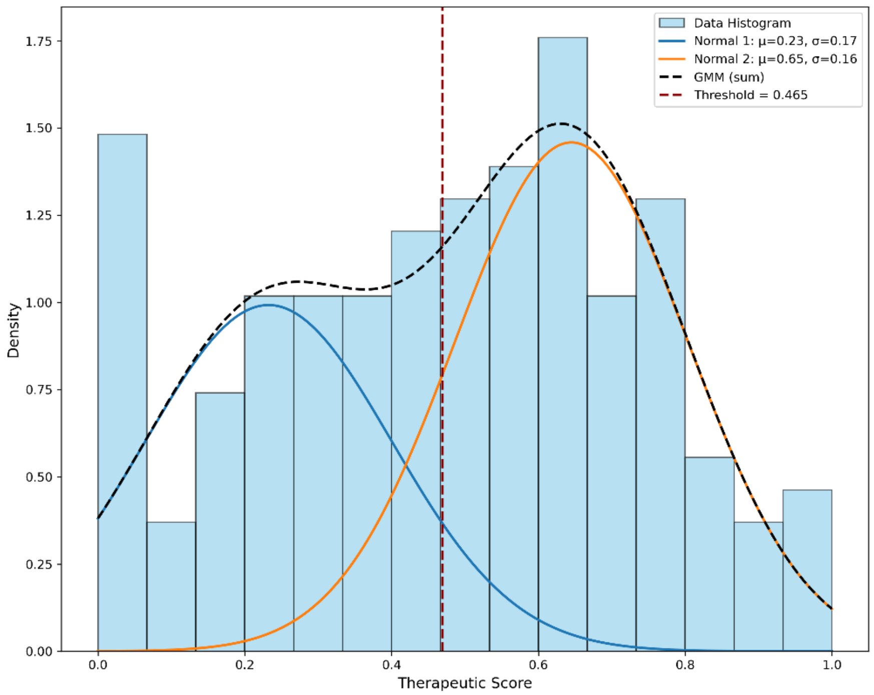


**Figure S1. Distribution of the reported therapeutic score.** The distribution was fitted with a two-component Gaussian Mixture Model. A classification threshold (red dashed line) was established at 0.465, a value near the intersection of the two component probability density functions.

To harmonize the mechanical properties measured by different approach, the storage modulus (G') was converted to elastic modulus (E) using the approximation E = 3G'.

The Changes in overall cytokines was derived by first normalizing the change for each reported cytokine against its control group and then averaging these normalized values.

### Multicollinearity Analysis

To ensure the quality of the data for training machine learning (ML) models, we examined the dataset for potential multicollinearity. A Pearson correlation matrix was generated to quantify the inter-correlations between the seven selected input features (**Figure S2**). As the heatmap shows, all pairwise correlation coefficients were low. The highest observed absolute correlation (0.3) was between "Materials" and "Delivery strategy". This widespread lack of strong correlations indicates that the features are sufficiently independent. This analysis validates the selection of these seven features for ML model training, as it confirms that multicollinearity would not be a confounding factor in the model's performance or its subsequent interpretation.

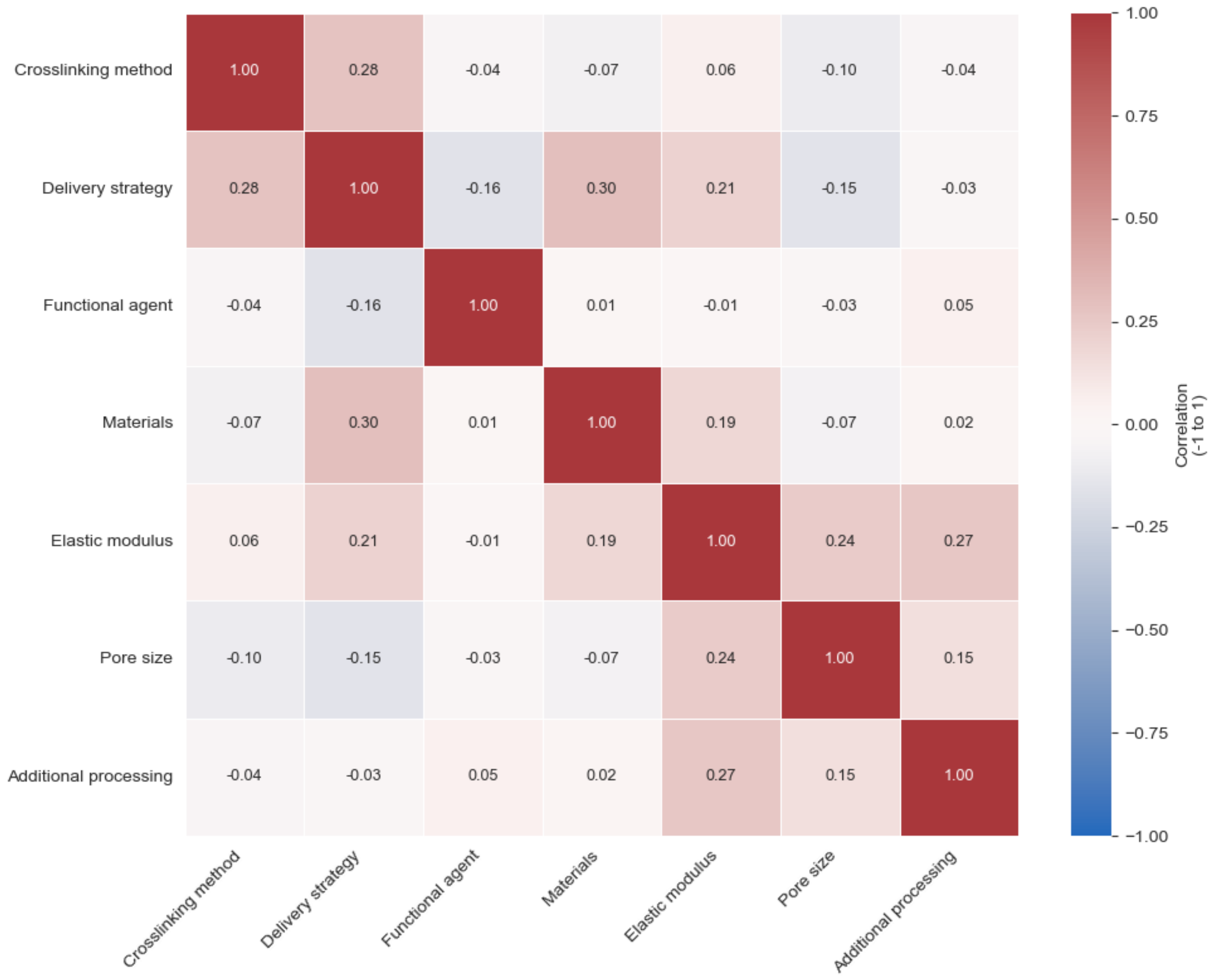


**Figure S2**. Correlation Matrix of Input Features. Heatmap displaying the Pearson correlation coefficients (r) between the seven harmonized features used for the machine learning analysis.

### External validation of the classification models

For the external validation, 79 valid entries were collected from papers published between October 1, 2024, and July 1, 2025. The trained models were then tested on this unseen data to examine their generalizability. SVM and Logistic Regression showed moderate performance, with accuracies of 0.63 (**Figure S3a**) and 0.61 (**Figure S3b**), respectively. The remaining models performed poorly, with XGBoost achieving 0.43 in accuracy (**Figure S3c**), KNN achieving 0.53 accuracy (**Figure S3d**) and Naïve Bayes 0.44 accuracy (**Figure S3e**). These models' performance was no better than random guessing.

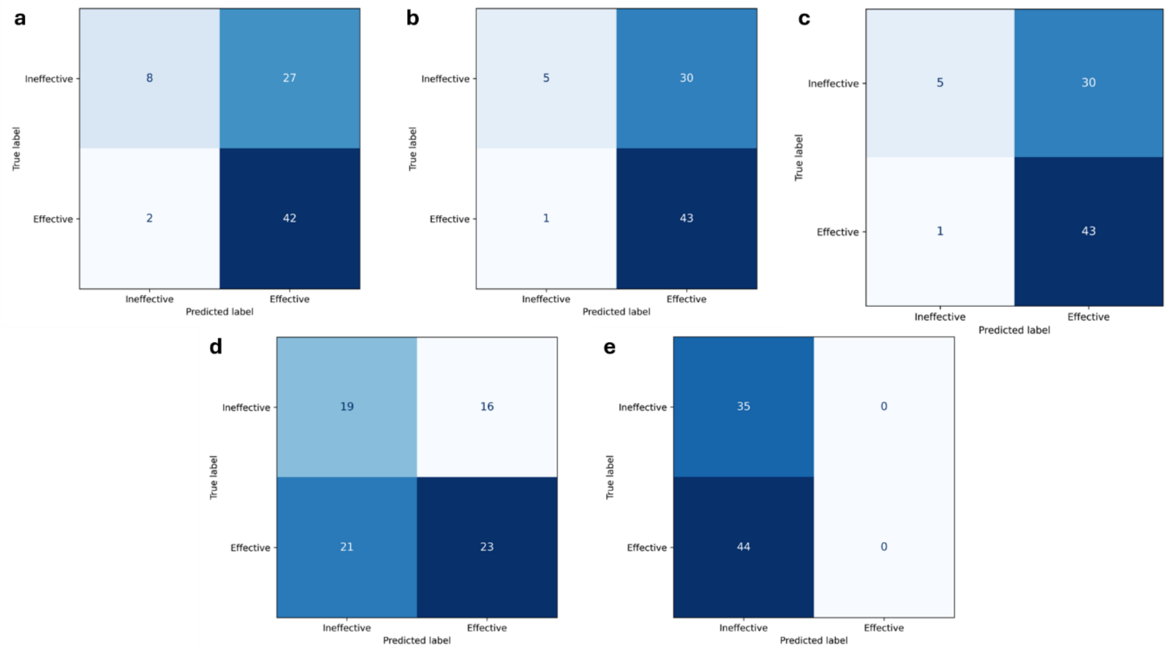


**Figure S3**. Confusion matrices for external validation on unseen data. Performance of the trained machine learning models on the prospective validation dataset (n=79 entries), which was curated from literature published after training models based on (a) SVM, (b) Logistic Regression, (c) XGBoost, (d) KNN, and (e) Naïve Bayes.

### SHAP analysis of the classification models

**Figure S4** shows the SHAP analysis on all five of the other classification models. The results across different models confirmed a consistent trend that “Functional agent” was ranked as the most important predictive feature across. “Materials” was also ranked as the second most important feature in three of the models. However, the analysis also revealed key differences in model complexity. The SHAP summaries for XGBoost (**Figure S4c**) and K-Nearest Neighbors (KNN) (**Figure S4d**) show a more distributed feature importance, where multiple factors (e.g., pore size, elastic modulus) clearly contribute to the prediction. This suggests these models captured more of the dataset’s complexity. In contrast, the simpler models appeared to underfit or over-focus. The Logistic Regression model (**Figure. S4b**) attributed almost all of its predictive power to the “Functional agent”, effectively ignoring other factors. Similarly, the Naïve Bayes model (**Figure S4e**) focused heavily on “Functional agent” and “Elastic modulus” while discounting the rest. The Support Vector Machine (SVM) (**Figure S4a**) showed a slightly better distribution but was still dominated by the top two features. This comparative analysis strengthens our confidence that “Functional agent” and “Materials” are the true dominant features, while also supporting our choice of a more complex model (i.e. Random Forest) that can capture the nuanced effects of secondary features.

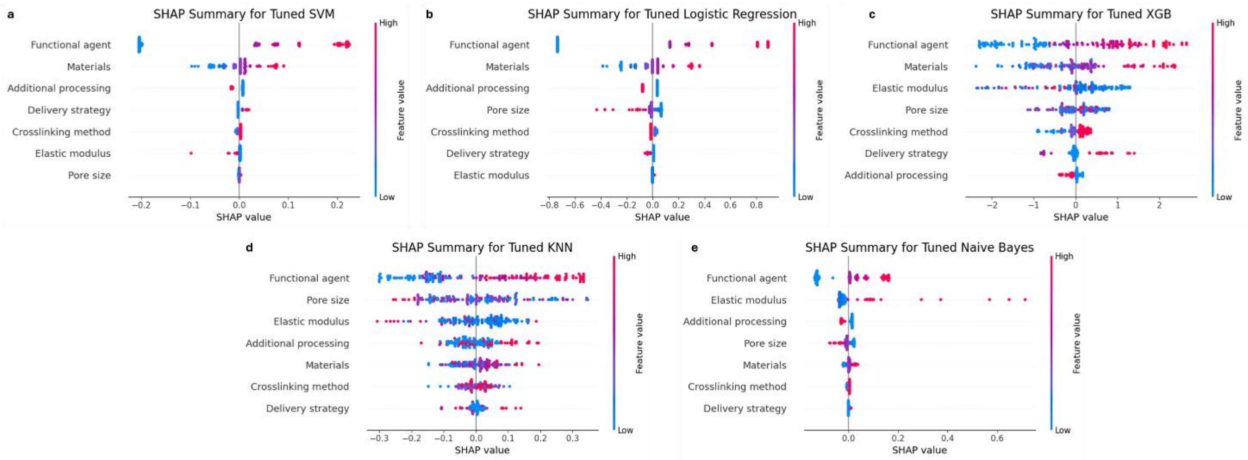


**Figure S4**. Comparative SHAP Summary Plots for All Classification Models. SHAP analysis was performed on the five additional machine learning models to assess feature importance consistency: (a) SVM, (b) Logistic Regression, (c) XGBoost, (d) KNN, and (e) Naïve Bayes.

#### Distribution of categorized materials and functional agents

For both the predictive ML and large language model (LLM) reasoning, raw material and agent names were classified into distinct, standardized categories. The distribution of these categories across the curated dataset is shown in **Figure S5**. The primary polymer materials (**Figure S5a**) are dominated by three classes: polysaccharide-based (32.2%), protein-based (29.8%), and synthetic polymers (29.5%), which collectively account for over 91% of all entries. The distribution of functional agents (**Figure S5b**) reveals that a significant portion of studies (37.3%) reported hydrogels with "No Functional Agent", relying instead on the intrinsic properties of the base material. This was followed by biological agents (17.3%), a category that includes cells, growth factors, and exosomes, and composite formulations (14.5%), which are defined as incorporating more than one type of functional agent. A detailed breakdown of the "Composite" category (**Figure S5c**) shows that the most common strategy (37.5%) is the combination of "Clinically Approved Drugs" and "Nanomaterials" followed by the second most common combination of "Clinically Approved Drugs", "Nanomaterials", and "Organic Compound". Notably, "Organic Compound" refers to the bioactive organic molecules that have not been clinically approved. This suggests a prevalent therapeutic approach where a drug is encapsulated within a nanocarrier to improve its delivery and efficacy.

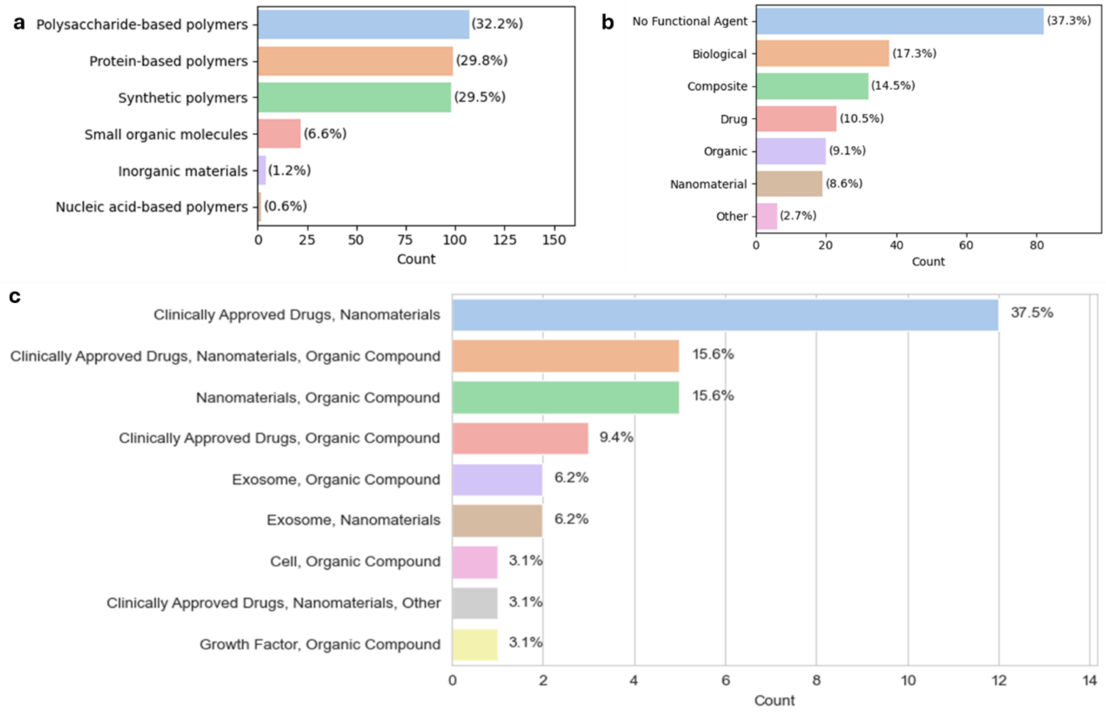


**Figure S5**. Distribution of categorized materials and functional agents used in hydrogels. Bar charts showing the frequency and percentage of (a) the categories of the polymer used, (b) the categorized functional agents. (c) Detailed breakdown of the "Composite" category from (b).
